# Gene-drug interactions identify genomic loci that enhance statin effectiveness in lowering LDL cholesterol

**DOI:** 10.64898/2026.02.04.703692

**Authors:** Brad Verhulst, Jennifer Harris, Amy M. Adams, Sarah E. Benstock, Carl W. Tong, Adam J. Case, John M. Hettema

## Abstract

Hyperlipidemia, and high low-density lipoprotein cholesterol (LDL-c) in particular, is a risk factor for cardiovascular disease, including atherosclerosis, myocardial infarction, and stroke. Nearly 200 million people worldwide take HMG-CoA reductase inhibitors, commonly known as statins, to lower their LDL-c. If statins interfere with the genetic pathways that endogenously increase the risk for hyperlipidemia, gene-statin interactions may identify genomic variants, and thereby individuals with those genotypes, that are particularly sensitive to these medications. We performed a series of genome-wide gene-statin interaction analyses in the UK Biobank for LDL-c and two related lipids: high-density lipoprotein cholesterol (HDL-c) and triglycerides (TG). We identified five genome-wide significant gene-statin interactions for LDL-c, two interactions for HDL-c, and four interactions for TG. Importantly, only the SNP-based heritability of LDL-c was reduced by statin use. Using data from All of Us, we replicated all five significant gene-statin interaction loci for LDL-c in the European-like ancestry sample, two loci in the Americas-like ancestry sample, and one locus in the African-like ancestry sample. We also identified fifteen loci that remained associated with LDL-c despite statin treatment, highlighting potential additional genetic targets for drug development, enhancement, and repurposing. These loci include gene-targets for the recently developed hyperlipidemia drug class (*PCSK9* inhibitors) validating our approach to finding new treatments. These results are an important step towards personalized medicine for patients with hyperlipidemia.

**Author Summary:** High cholesterol raises the risk of heart attacks and strokes and nearly 200 million people worldwide take statins to lower it. While statins work for nearly everyone, they work better for some people than others. We examined how genetic differences enhance the effectiveness of statin medication as a step toward enhancing personalized medicine for those with high cholesterol. By analyzing genetic and health data from about 390,000 people, we found that statins primarily disrupt the link between genes and LDL or “bad” cholesterol levels. Across the genome, statins reduce the impact of genes on LDL-c levels, but not other blood lipids like HDL-c or triglycerides. For LDL-c specifically, we found five regions where genetic differences increase the effectiveness of statins. People with the protective genetic variants are expected to see greater cholesterol reduction with statin use. We confirmed four of the five findings in European-like ancestry samples, with partial replication in Americas-like and African-like ancestry samples (two variants and one variant, respectively). We also found 15 genomic regions where cholesterol stays high despite statin treatment. These genomic regions could be targets for new or enhanced cholesterol medications. In fact, a newer drug class targets one of these regions supporting our approach to finding new treatments.

## Introduction

Hyperlipidemia, characterized by aberrant lipid concentrations in the blood, increases the risk for atherosclerosis, myocardial infarction, and stroke. High levels of low-density lipoprotein cholesterol (LDL-c) appear to play the largest causal role in this process^1^, but high levels of triglycerides (TG) and low levels of high-density lipoprotein cholesterol (HDL-c) may also contribute to adverse cardiovascular outcomes. Lipid levels have a strong genetic component, which has two implications. First, because lipid levels are, in part, a function of genetic processes, targeting the genetic mechanisms that endogenously affect lipid levels may interfere with these genetic pathways and reduce the association between genes and lipid profiles. Second, because everyone has a unique genotype, pharmaceutical interventions may be more effective for certain people. Interactions between statins and single nucleotide polymorphisms (SNPs) within gene regions can amplify (or reduce) drug effectiveness. Such gene-drug interactions highlight potential genomic mechanisms that regulate lipid metabolism and homeostasis, identify genetic markers for individuals who may benefit the most (or least) from statin treatment, and lay the foundation to develop proactive and personalized solutions to reduce cardiovascular disease (CVD) morbidity and mortality^2,3^.

The primary treatment for hyperlipidemia is statins, which reduce LDL-c levels in almost everyone^4–8^, but approximately 10% of patients experience side effects such as myopathy or hepatic and renal inflammation^9^ potentially due to genetic variation^10,11^. While people who experience adverse events from statins are likely to stop or change treatments^12,13^, others may unknowingly benefit more than average. With an estimated 145–200 million statin users globally^14^ and utilization increasing approximately 25% between 2015 and 2020^15^, even modest improvements in treatment optimization could have substantial clinical impact. Existing pharmacogenomic research for statins has identified 8 loci that increase the effectiveness of statins for lowering LDL-c^13,16,17^: *SORT1*/*CELSR2*/*PSRC1*, *ABCG2*, *KIF6*, *LPA*, *CYP2C9*, *SLCO1B1*, *APOE* and *LDLR*. Notably, the evidence for many of these pharmacogenomic loci is based on a limited number of genomic variants (often one SNP) despite very high levels of linkage disequilibrium (LD) in the regions. Furthermore, while these loci are associated with lipid levels, they are not the most significant genetic associations in a recent, large-scale, multi-ancestry lipid GWAS^18^. This implies that the genetic architecture of statin responsiveness and serum lipid levels is moderately distinct.

Our premise is that genetic factors increase an individual’s endogenous risk for hyperlipidemia and statins reduce this risk by interfering with these genetic pathways. This suggests three hypotheses. First, if statins interfere with the genetic pathways that culminate in hyperlipidemia, then the heritability, or the variation accounted for by genetic factors, of lipids should be lower in individuals taking statins. Second, identifying SNPs implicated in gene-drug interactions can highlight biological mechanisms and identify individuals who may benefit the most from statin use. Third, genetic associations for individuals already taking statins can highlight putative gene-targets for drug improvement, repurposing, and novel development. The preponderance of causal evidence linking lipids to CVD focuses on LDL-c^1,19^, but given correlations between LDL-c, HDL-c, and TG it is instructive to explore the possibility of statin effects on these related outcomes. As causal evidence is stronger for LDL-c, pharmacogenomic hypotheses apply more directly to LDL-c than to either TG or HDL-c. In this study, we test these hypotheses by conducting genome-wide gene-statin interaction analyses for LDL-c, HDL-c, and TG with data from the European ancestry subset of the UK Biobank (see Methods). To replicate the findings, we conducted targeted follow-up analyses in the European-like, African-like, and Americas-like ancestry subsamples of the All of Us (AoU) data^20^ based on the genome-wide significant gene-statin interactions in the UKB discovery analyses.

## Results

### Sample Characteristics

The means and standard deviations of the three blood lipid levels, the proportion of people taking a statin, and other relevant demographic information for the UK Biobank (UKB) sample are presented in **Table 1**. Detailed drug codes are described in **Supplementary Table 1**. Corresponding information for the All of Us (AoU) sample is presented in **Table 2**. Across both samples, statin users are older, more likely to be male, have lower levels of SES (based on income, TDI and/or educational attainment), and are more likely to be current or former smokers. In the UKB sample, statin users have higher BMIs, and slightly higher systolic (but not diastolic) blood pressure. Nevertheless, we do not covary for these cardiovascular, behavioral, and sociodemographic variables in the GxE GWAS analyses to avoid introducing collider bias into the estimated genetic associations^21,22^.

**Table 1:** Descriptive statistics for the key variables in the GxE GWAS analysis of Low Density Lipoprotein Cholesterol (LDL-c), High Density Lipoprotein Cholesterol (HDL-c) and Triglycerides (TG).

|  | Full Sample | Statin Users** | Statin Non-Users |
| --- | --- | --- | --- |
| **Sample Size (%)** | 387,420 | 63,517 (16.3) | 323,923 (83.7) |
| Atorvastatin (%) |  | 14,378 (22.6) |  |
| Fluvastatin (%) |  | 197 (0.3) |  |
| Pravastatin (%) |  | 1,868 (2.9) |  |
| Rosuvastatin (%) |  | 2,623 (4.1) |  |
| Simvastatin (%) |  | 44,501 (70.0) |  |
| **Lipids (mg/dL)\*** |  |  |  |
| LDL (sd) | 137.94 (33.58) | 107.01 (26.04) | 143.97 (31.50) |
| HDL (sd) | 56.22 (14.82) | 50.61 (13.70) | 57.31 (14.78) |
| TG (sd) | 155.05 (90.59) | 172.65 (96.21) | 151.62 (89.05) |
| **Covariates** |  |  |  |
| Age (sd) | 57.26 (8.01) | 62.05 (5.97) | 56.33 (8.02) |
| Females (%) | 210,169 (54.0) | 24,474 (38.5) | 185,695 (57.0) |
| **Cardiovascular Outcomes** |  |  |  |
| BMI (sd) | 27.38 (4.77) | 29.37 (5.01) | 26.99 (4.63) |
| Systolic BP (sd) | 137.99 (18.65) | 142.11 (17.97) | 137.19 (18.67) |
| Diastolic BP (sd) | 82.23 (10.14) | 82.12 (9.91) | 82.25 (10.18) |
| **Behavioral and Demographic Outcomes** |  |  |  |
| Alcohol Use Frequency (sd) | 3.14 (1.49) | 3.04 (1.59) | 3.16 (1.46) |
| Townsend Deprivation Index (sd) | -1.45 (3.00) | -1.13 (3.18) | -1.52 (2.96) |
| Tobacco Smoking Status |  |  |  |
| Current Smoker (%) | 40,988 (10.6) | 7,226 (11.4) | 33,762 (10.4) |
| Former Smoker (%) | 137,932 (35.5) | 28,788 (45.6) | 109,144 (33.6) |
| Never Smoked (%) | 209,181 (53.9) | 27,184 (43.0) | 181,997 (56.0) |
| Educational Attainment |  |  |  |
| A Levels (%) | 44,429 (11.5) | 5,776 (9.2) | 38,653 (12.0) |
| Degree (%) | 127,951 (33.2) | 15,319 (24.4) | 112,632 (34.9) |
| GCSE (%) | 128,408 (33.3) | 20,182 (32.2) | 108,226 (33.5) |
| None of the Above (%) | 64,933 (16.8) | 17,557 (28.0) | 47,376 (14.7) |
| Other (%) | 19,946 (5.2) | 3,859 (6.1) | 16,087 (5.0) |

Note: Participants were excluded from the sample if they reported ONLY taking a non-statin cholesterol lowering medication (e.g., Ezetimibe). Individuals who reported ONLY taking

**Table 2:** Descriptive statistics for the key variables in the GxE GWAS analysis of Low Density Lipoprotein Cholesterol (LDL-c) in the All of Us Samples.

|  | AFR-like Ancestry Group |  |  | AMR-like Ancestry Group |  |  | EUR-like Ancestry Group |  |  |
| --- | --- | --- | --- | --- | --- | --- | --- | --- | --- |
|  | All | Statin | Non-Statin | All | Statin | Non-Statin | All | Statin | Non-Statin |
| LDL-c (mg/dL)* | 100.9 (31.1) | 99.4 (33.8) | 101.6 (29.9) | 102.9 (29.9) | 97.6 (33.0) | 103.9 (29.1) | 102.7 (29.3) | 91.3 (28.9) | 106.8 (28.4) |
| Any Statin (%) | 18,719 | 5,562 (29.7) | 13,157 (70.3) | 16,074 | 2,743 (17.1) | 13,331 (82.9) | 69,123 | 18,184 (26.3) | 50,939 (73.7) |
| Atorvastatin (%) |  | 4,992 (89.8) |  |  | 2,483 (90.5) |  |  | 15,243 (83.8) |  |
| Cerivastatin (%) |  | <20 (0.0) |  |  | <20 (0.0) |  |  | 23 (0.1) |  |
| Fluvastatin (%) |  | 863 (15.5) |  |  | 495 (18.0) |  |  | 3,410 (18.8) |  |
| Lovastatin (%) |  | 424 (7.6) |  |  | 401 (14.6) |  |  | 1,771 (9.7) |  |
| Pitavastatin (%) |  | 30 (0.5) |  |  | 13 (0.5) |  |  | 192 (1.1) |  |
| Pravastatin (%) |  | 1,928 (34.7) |  |  | 785 (28.6) |  |  | 5,658 (31.1) |  |
| Rosuvastatin (%) |  | 2,049 (36.8) |  |  | 908 (33.1) |  |  | 7,573 (41.6) |  |
| Simvastatin (%) |  | 3,154 (56.7) |  |  | 1,654 (60.3) |  |  | 11,120 (61.2) |  |
| **Covariates** |  |  |  |  |  |  |  |  |  |
| Age (sd) | 57.8 (14.6) | 67.9 (10.0) | 53.5 (14.1) | 52.9 (15.1) | 68.2 (10.8) | 49.8 (14.0) | 61.9 (16.5) | 74.1 (10.2) | 57.6 (16.1) |
| Females (%) | 12,508 (67.9) | 3,387 (62.2) | 9,121 (70.3) | 11,099 (69.6) | 1,611 (59.4) | 9,488 (71.7) | 42,268 (61.6) | 8,149 (45.4) | 34,119 (67.4) |
| **Behavioral and Demographic Outcomes** |  |  |  |  |  |  |  |  |  |
| Alcohol Use Frequency (sd)** | 1.40 (1.14) | 1.17 (1.13) | 1.50 (1.13) | 1.30 (1.07) | 1.00 (1.06) | 1.35 (1.07) | 1.89 (1.31) | 1.75 (1.38) | 1.94 (1.29) |
| Tobacco Smoking Status |  |  |  |  |  |  |  |  |  |
| Current Smoker (%) | 4,233 (23.3) | 1,283 (23.6) | 2,950 (23.1) | 1,640 (10.5) | 257 (9.5) | 1,383 (10.6) | 5,982 (8.7) | 1,514 (8.4) | 4,378 (8.8) |
| Former Smoker (%) | 3,199 (17.6) | 1,447 (26.6) | 1,752 (13.7) | 2,814 (18.0) | 697 (25.7) | 2,117 (16.4) | 21,551 (31.9) | 7,706 (42.9) | 13,845 (27.8) |
| Never Smoked (%) | 10,754 (59.1) | 2,667 (49.0) | 8,087 (63.2) | 11,176 (71.5) | 1,731 (63.9) | 9,445 (73.0) | 40,194 (59.4) | 8,613 (48.0) | 31,581 (63.4) |
| Income (sd)*** | 39,114 (45,000) | 31,876 (38,304) | 42,054 (47,135) | 51,478 (53,659) | 36,245 (45,607) | 54,405 (54,589) | 97,981 (68,012) | 84,902 (63,599) | 102,497 (68,897) |
| Educational Attainment |  |  |  |  |  |  |  |  |  |
| Advanced Degree (%) | 1,920 (10.3) | 429 (7.7) | 1,491 (11.3) | 1,526 (9.5) | 141 (5.1) | 1,385 (10.4) | 22,526 (32.6) | 5,290 (29.1) | 17,236 (33.8) |
| College Degree (%) | 2,971 (15.9) | 703 (12.6) | 2,268 (17.2) | 2,640 (16.4) | 289 (10.5) | 2,351 (17.6) | 20,418 (29.5) | 4,490 (24.7) | 15,928 (33.8) |
| Some College (%) | 5,892 (31.5) | 1,735 (31.2) | 4,157 (31.6) | 3,932 (24.5) | 554 (20.2) | 3,378 (25.3) | 16,533 (23.9) | 5,043 (27.7) | 11,490 (22.6) |
| High School (%) | 5,159 (27.6) | 1,591 (28.6) | 3,568 (27.1) | 3,473 (21.6) | 573 (20.9) | 2,900 (21.8) | 7,587 (11.0) | 2,699 (14.8) | 4,888 (9.6) |
| No High School (%) | 2,096 (11.2) | 855 (15.4) | 1,241 (9.4) | 4,056 (25.2) | 1,104 (40.2) | 2,952 (22.1) | 1,312 (1.9) | 414 (2.3) | 898 (1.8) |

Note: As the All of Us data are potentially longitudinal, participants could have multiple measures for each variable, including times when they took and did not take statins. If a person took a statin at least once, they were considered a statin user. To include as much data as possible, the average LDL-c across multiple measures were averaged within participants. For statin users, only LDL-c concentrations that were measured during the prescription window were included in the mean. Observations were excluded if the individual reported ONLY taking a non-statin cholesterol lowering medication (e.g., Ezetimibe). Individuals who reported ONLY taking vitamins or non-pharmacological herbal remedies for statin use were included in the analyses and considered non-statin users (e.g., Fish Oil Supplements). For “any statin”, the percentages reflect any statin vs no statin. For the individual statins, the percentages reflect if the specific statin was prescribed for the respondent at least once.
\* Lipid concentrations are reported in mmol/l. If concentrations were reported in alternative formats, we converted the values from mmol/l to mg/dl by multiplying the observed values for LDL-c 38.67. These transformations have no impact on the conclusions. \*\*Statin Users - Statin use was coded as a binary variable based upon self-reported use vs non-use of any statin. The coding did not integrate dosage information or adherence to treatment regimen.
\*\* Alcohol use frequency was measured on an ordinal scale (0 = Never drinks; 1 = Monthly or less; 2 = 2-4 Drinks per month; 3 = 2-3 drinks per week; 4 = more than 4 per week).
\*\*\* Income is reported in dollars.

### Phenotypic Correlations

LDL-c, HDL-c, and TG are moderately correlated. In the UKB (details in **Methods**), statin use induces a small phenotypic correlation between LDL-c and HDL-c (r_no statin_ =0.01, r_statin_ = 0.16) and modestly reduce the phenotypic correlation between LDL-c and TG (r_no statin_ =0.29, r_statin_ = 0.24). Thus, statin use alters the pattern of phenotypic correlations between the lipids, potentially by interfering with the genetic mechanisms regulating LDL-c.

### Heritability and Genetic Correlations

We test whether statins reduce the SNP-based heritability (h^2^_SNP_) of each lipid. We used the summary statistics from the gene-statin interaction analyses to calculate genetic marginal effects for statin users and non-users for each lipid^23^ (**See Methods**). We used the marginal genetic effects to estimate h^2^_SNP_ of each lipid using LD score regression (LDSC)^24^ depending on statin use^25^. Statin use selectively reduced h^2^_SNP_ for LDL-c (non-users: 0.14, SE = 0.01; users: 0.09, SE = 0.01) while h^2^_SNP_ of HDL-c and TG remained unchanged (**Figure 1; Supplementary Table 2**). Notably, as lipid levels had h^2^_SNP_ of approximately 10-20%, which is notably smaller than family-based heritability estimates^26,27^, a sizeable portion of the variation in lipids may be explained by rare genetic variation or environmental factors such as diet or exercise behaviors. The specific decrease in h^2^_SNP_ for LDL-c is consistent with clinical observations where statins reduce LDL-c levels and minimally affect TG and HDL-c^28^. Further, SNP-based genetic correlations (r_G_) between statin users and non-users for LDL-c were significantly less than one (r_G_ = 0.79, p_rG<1_ = 0.016), implying that somewhat different sets of genetic factors contribute to the genetic architecture of LDL-c depending on statin use. The r_G_ between statin users and non-users for HDL-c and TG are not different from one (HDL-c: r_G_ = 0.95, p_rG<1_ = 0.55; TG: r_G_ = 0.99, p_rG<1_ = 0.89). The observation that LDL-c h^2^_SNP_ decreases with statin use and the r_G_ for LDL-c between statin users and non-users is less than unity suggests statin use disrupts specific genetic pathways affecting LDL-c while circumventing the genetic mechanisms that influence HDL-c and TG.

**Figure 1:**
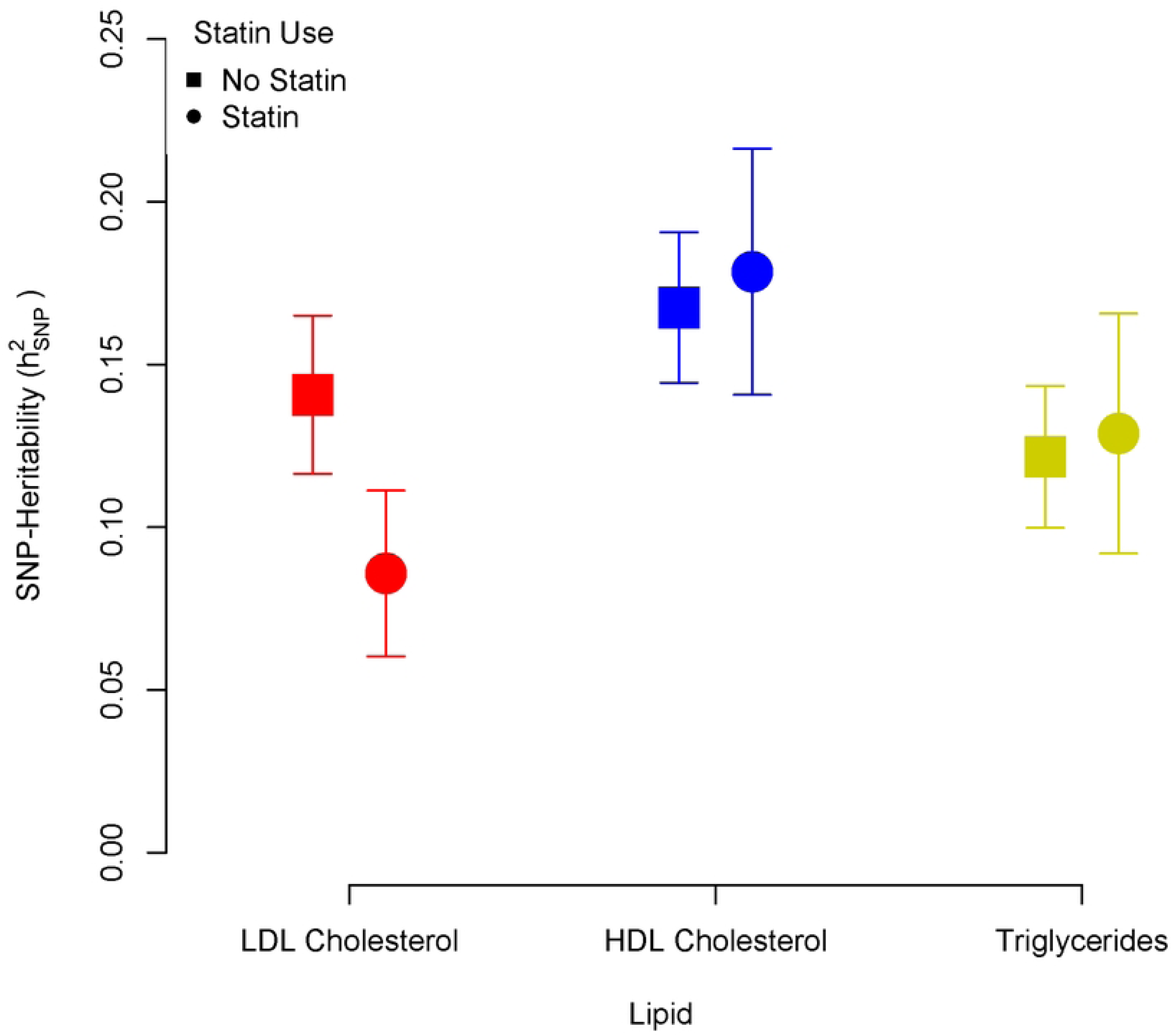
SNP-Heritability (h^2^_SNP_) of LDL Cholesterol (red), HDL Cholesterol (blue) and Triglyceride (yellow) levels as a function of statin use.

### SNP-statin Interactions in LDL-c, HDL-c and TG in the UK Biobank

Our discovery analysis identified five genome wide significant interactions between SNPs within a genomic loci and statin use for LDL-c (**Table 3***; rs934197; rs1535; rs17242381; rs10401969*; and *rs72654473*), two interactions for HDL-c *(rs247616*; and *rs429358*), and four interactions for TG (*rs55730499; rs102275; rs7118999*; and *rs7412*). Interestingly, the SNP-statin interactions in the *APOE*/*PVRL2* region for LDL-c, HDL-c, and TG analyses identified different lead SNPs for each lipid (rs72654473, rs429358, and rs7412, respectively; **Figure 2)**, despite the phenotypic correlations and high LD in the region. Thus, while we do not assume the variants we identified are causal, it is clear this region has a strong, association with broad lipid regulation that is altered by statin use. Specifically, in the *APOE*/*PVRL2* region, statins reduce (but do not eliminate) the marginal genetic effects of the SNPs on LDL-c and HDL-c, with the pattern of effects being stronger and more consistent for LDL-c. For TG, statins increase the effect size of the *APOE/PVRL2* region. This pattern of results is consistent with the observation that taking statins may induce a small, positive phenotypic correlation between LDL-c and HDL-c and reduce the correlation between LDL-c and TG. The global pattern of reduced marginal genetic effects is consistent across the other genome-wide significant interaction loci (p < 5×10^-8^) for LDL-c and HDL-c, but variable for TG depending on the genomic loci. Overall, the patterns of SNP-statin interactions for the three lipids are relatively distinct (see **Supplementary Figures 4-14** for regional plots of additional loci).

**Figure 2:**
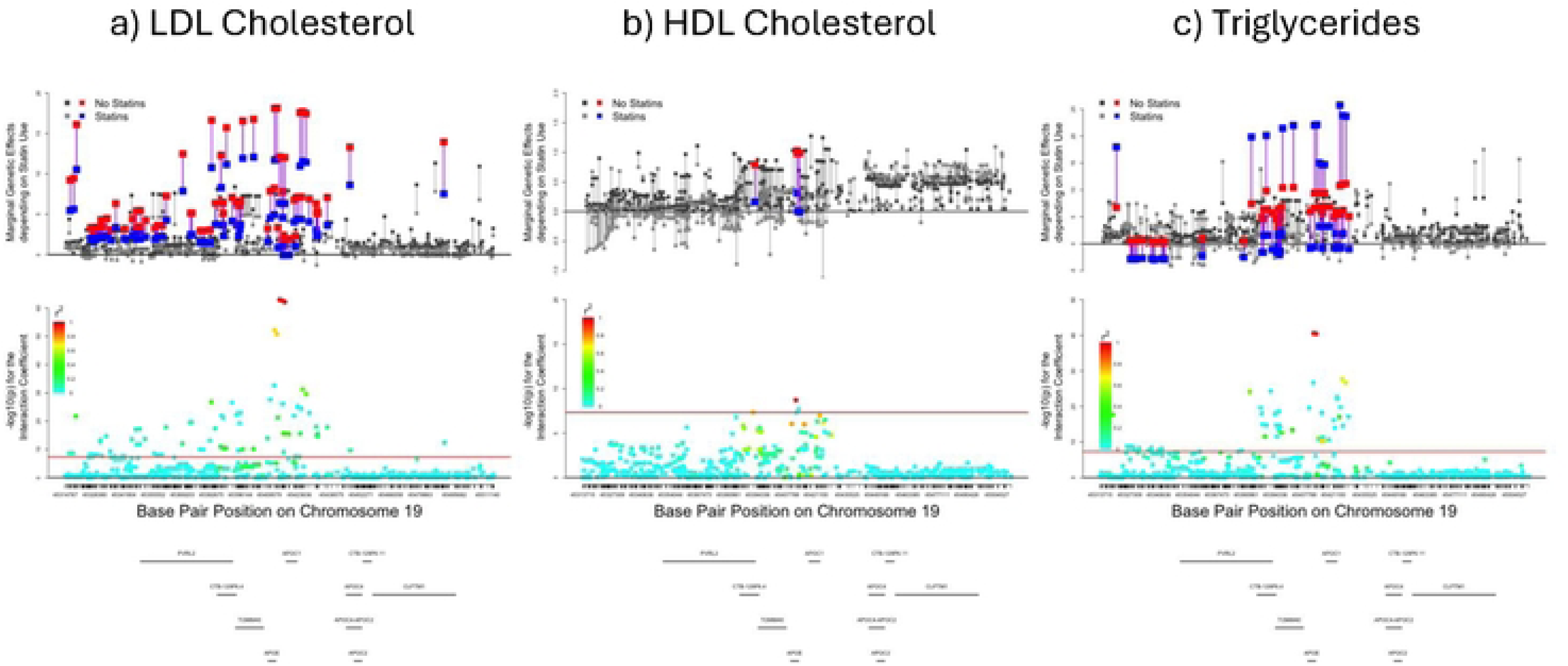
Locus plot of the genetic marginal effects for statin users and non-users and the regional Manhattan plots of the gene-statin interaction coefficient for the APOE region in LDL Cholesterol, HDL Cholesterol and Triglycerides.

**Table 3:** Results for the genome-wide significant gene-statin interaction index variants identified in the GxE GWAS analysis of LDL Cholesterol, HDL Cholesterol and Triglycerides in 387,420 individuals.

| Index Variant | Chr | BP | A1 | A2 | Main Effect ($\beta_1$) | | | | Interaction Effect ($\beta_3$) | | | | Nearest Gene | eQTL Gene | eQTL min p-value | Number of Indep Sig SNPs |
| --- | --- | --- | --- | --- | --- | --- | --- | --- | --- | --- | --- | --- | --- | --- | --- | --- |
| | | | | | $\beta_1$ | SE$_1$ | Z$_1$ | P$_1$ | $\beta_3$ | SE$_3$ | Z$_3$ | P$_3$ | | | | |
| LDL Cholesterol |  |  |  |  |  |  |  |  |  |  |  |  |  |  |  |  |
| rs934197 | 2 | 21267461 | A | G | 3.67 | 0.08 | 46.19 | <5 e-310 | -1.35 | 0.22 | -6.14 | 8.31 e-10 | APOB | APOB | 4.92 e-29 | 3 |
| rs1535 | 11 | 61597972 | A | G | 1.54 | 0.08 | 19.36 | 1.53 e-83 | -1.72 | 0.22 | -7.70 | 1.40 e-14 | FADS2 | FADS2 | <5 e-310 | 2 |
| rs17242381 | 19 | 11206729 | T | C | 6.21 | 0.12 | 53.29 | <5 e-310 | -3.17 | 0.34 | -9.42 | 4.73 e-21 | LDLR | SMARCA4 | 2.63 e-68 | 6 |
| rs10401969 | 19 | 19407718 | T | C | 4.42 | 0.14 | 31.24 | 3.35 e-214 | -2.53 | 0.41 | -6.19 | 6.19 e-10 | SUGP1 | MAU2 | <5 e-310 | 1 |
| rs72654473 | 19 | 45414399 | C | A | 12.11 | 0.11 | 105.34 | <5 e-310 | -5.75 | 0.34 | -16.84 | 1.20 e-63 | APOE | PVRL2 | <5 e-310 | 41 |
| HDL Cholesterol |  |  |  |  |  |  |  |  |  |  |  |  |  |  |  |  |
| rs247616 | 16 | 56989590 | C | T | 3.52 | 0.04 | 97.31 | 0.00 e+00 | -0.58 | 0.09 | -6.18 | 6.36 e-10 | CETP | CETP | 1.10 e-58 | 2 |
| rs429358 | 19 | 45411941 | C | T | -1.03 | 0.05 | -20.57 | 4.67 e-94 | 0.71 | 0.12 | 6.00 | 1.96 e-09 | APOE | PVRL2 | 1.66 e-28 | 2 |
| Triglycerides |  |  |  |  |  |  |  |  |  |  |  |  |  |  |  |  |
| rs55730499 | 6 | 161005610 | C | T | -2.78 | 0.43 | -6.41 | 1.48 e-10 | -5.81 | 0.93 | -6.21 | 5.23 e-10 | LPA | LPA | 5.83 e-05 | 2 |
| rs102275 | 11 | 61557803 | T | C | 4.10 | 0.24 | 17.32 | 3.18 e-67 | 4.20 | 0.53 | 7.91 | 2.48 e-15 | FADS2 | FADS2 | <5 e-310 | 2 |
| rs7118999 | 11 | 116645275 | C | T | -11.91 | 0.25 | -47.27 | <5 e-310 | -3.28 | 0.56 | -5.85 | 5.04 e-09 | APOA4 | BUD13 | 1.39 e-27 | 1 |
| rs7412 | 19 | 45412079 | C | T | 9.38 | 0.39 | 24.19 | 2.86 e-129 | 12.63 | 0.94 | 13.46 | 2.68 e-41 | APOE | PVRL2 | 3.14 e-282 | 14 |

### Variant-level differences in the genetic marginal effects for statin users versus non-users

We present circular Manhattan plots of -log_10_ p-values of the marginal genetic effects in the no statin and statin groups in **Figures 3a and 3b**. We highlight genomic regions with significant SNP-statin interactions across lipids (standard Manhattan and QQ plots of all the marginal genetic effects are presented in **Supplementary Figures 1-3).** A complete list of the lead SNPs for the genome-wide significant marginal genetic effects for statin users and non-users in each lipid is presented in **Supplementary Tables 3-5**. The marginal genetic effects can be interpreted in the same way as stratified GWAS associations for a specific level of the moderator (i.e., genetic associations for non-statin or statin users). The red, blue, and yellow bands around the Manhattan plots in **Figure 3** denote genome-wide significant associations for LDL-c, HDL-c, and TG, respectively. The robust polygenic architecture of unmedicated respondents for all three lipids is evident in the myriad genome-wide significant associations in **Figure 3a**. The large number of genome-wide risk loci for non-statin users (LDL-c = 247, HDL-c = 234, TG = 139) reflects the central role genetic factors play in endogenous lipid regulation and metabolism^29^. The results for non-statin users reflect the findings from the literature^18^. Compared with the non-statin group, there is a muted genetic architecture for statin users as seen in **Figure 3b** (number of genome-wide significant risk loci for statin users: LDL-c = 15, HDL-c = 38, TG = 82). However, the differences in the magnitude of the p-values may reflect two factors. First, there is a large asymmetry in the sample sizes between statin users and non-users (N_no statins_ = 325,923 vs N_statins_ = 63,517) which reduces the statistical power and significance for the analyses of statin users. Second, statins may directly reduce the magnitudes of the genetic associations. In the current analyses, statin use reduces the magnitude of the genetic associations for LDL-c, while the difference in the magnitudes of the genetic associations for HDL-c and TG for most loci is negligible. Thus, the interpretation of the differences between the results for statin and non-statin users for LDL-c are consistent with SNP-statin interactions, while the interpretation of the results for HDL-c and TG is more consistent with differences in sample size.

**Figure 3:**
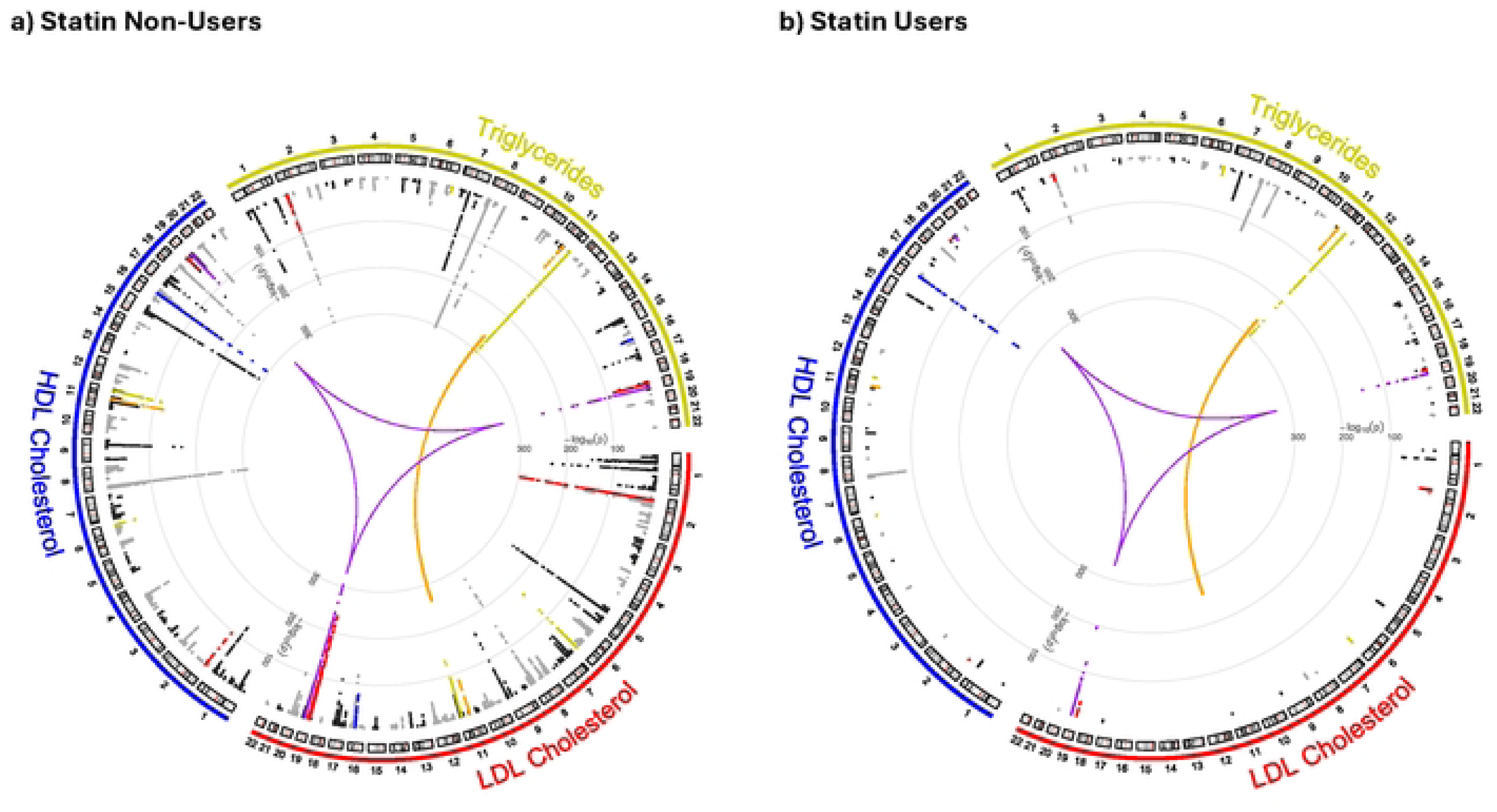
Circular Manhattan Plots of p-values for the marginal effects with significant interactions for LDL (red), HDL (blue), and TG (yellow) for people who do not take statins (a) and who do take statins (b).

### Integrating Pharmaceutical and Genomic Results

We integrated data from the LDL-c genome-wide gene-statin interaction analysis with the pharmGKB.org database^30,31^ (see **Supplementary Table 6** for the list of statin gene-targets) to address two distinct research questions: which interaction loci from our analysis correspond with existing evidence of pharmacogenomic sensitivity to medication (left edge of the triad of **Figure 4**), and are any of the loci that remain significant for statin users sensitive to existing medications that may highlight avenues for future pharmacological refinement or development (right edge of the triad of **Figure 4**). We present existing pharmacogenomic evidence from pharmGKB in the bottom of the triad of **Figure 4**. The pharmacogenomic analyses were restricted to gene-drug relationships with high (1) or moderate (2) levels of evidence for variant and clinical annotations, or “Informative PGx” or “Actionable PGx” levels of evidence for drug label annotations.

**Figure 4:**
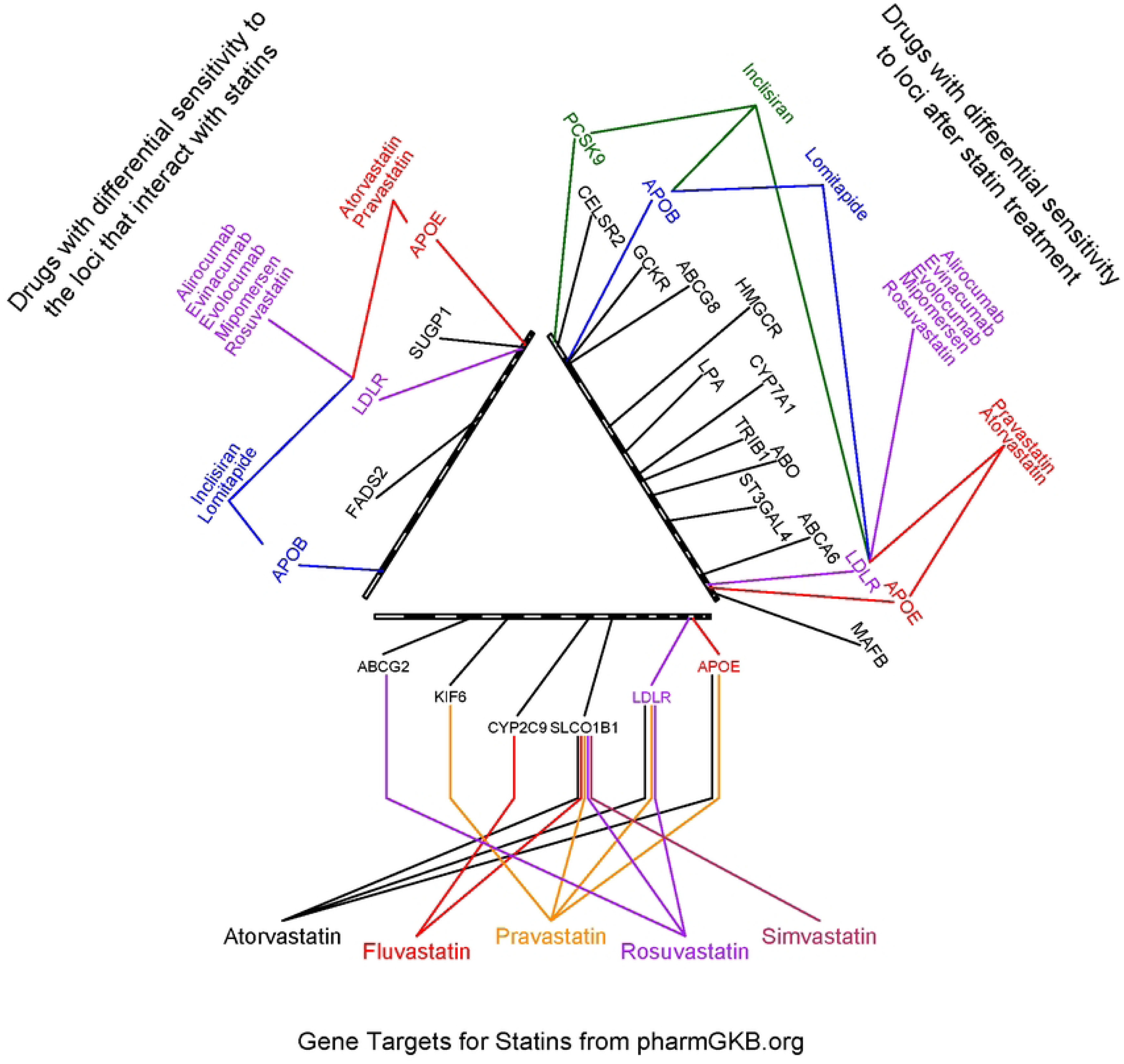
Pharmacogenomic analysis of statins.

The bottom of the triad of **Figure 4** lists gene targets and chromosome positions for the five statins included in our analysis. Existing evidence from pharmGKB emphasizes the centrality of *LDLR*, *APOE*, and *SLCO1B1* for individual statins. Notably, in addition to *APOE* and *SLCO1B1*, two additional loci have been identified: *SORT1*/*CELSR2*/*PSRC1* on chromosome 1 and *LPA* on chromosome 6^16^. These results provide the basis for existing expectations for empirically testing the influence of statins on LDL-c in the general population. Importantly, existing evidence suggests that each of the five statins is sensitive to *SLCO1B1* (which we did not replicate) but differed in their sensitivity to other lipid transport protein genes.

The top left of the triad of **Figure 4** presents the significant interaction loci for LDL-c with previously identified drug-gene targets to highlight pharmacodynamic mechanisms of statins. Two of the five genome-wide significant interaction loci from our analyses, *LDLR*/*SMARCA4* and *APOE*/*PVRL2*, have been previously identified as pharmacogenomic loci for statins, and *APOB* is sensitive to lomitapide and inclisiran which are non-statin medications used to lower LDL-c. In addition, we identified two novel pharmacogenomic loci: *FADS2* and *SUGP1*/*MAU2*. While both have previously been associated with LDL-c levels^18^, this is the first time they have been shown to interact with statins to further lower LDL-c. Notably, we did not detect interactions for *HMGCR* or *LPA* despite HMGCR encoding the purported molecular target of statins. This null finding is consistent with prior pharmacogenomic findings and may reflect the mechanism of statin action^16^. As competitive inhibitors that bind directly to the HMG-CoA reductase enzyme, statins may override genetic variation in HMGCR expression or activity at the protein level. In contrast, the loci we identified likely affect parallel pathways—cholesterol uptake, transport, and clearance—where genetic variation modulates how effectively statins reduce circulating LDL-c. For *LPA*, the lack of interaction with LDL-c is consistent with evidence that statins do not substantially lower lipoprotein(a) levels; notably, we did observe a significant *LPA*-statin interaction for TG^32^ suggesting an alternative potential mechanism.

The top right of the triad of **Figure 4** identifies 15 genetic associations with LDL-c that remained significant despite statin use. These residual genetic associations suggest gene targets for 1) drug improvement, 2) drug repurposing, and 3) novel drug development. The most straightforward implication of this analysis is for drug improvement. Specifically, while *LDLR*/*SMARCA4* (*rs17242381*) and *APOE*/*PVRL2* (*rs72654473*) increase the effectiveness of statins for individuals with the protective alleles, they remain significant after taking statins. This suggests that while currently available statins are effective, drug refinement that more precisely targets these loci could further reduce LDL-c. In addition, these residual genetic associations may highlight gene targets for novel drug development. Finding a residual genetic association for *PCSK9* (*rs1159147*) exemplifies the potential for novel drug discovery. *PCSK9* is the target gene of an emerging class of medications (*PCSK9* inhibitors) used to lower LDL-c that work on a different set of biological pathways than statins. Because *PCSK9* inhibitors were rarely prescribed to patients in our data, they were excluded from the analysis. Accordingly, our results “post-dict” the effectiveness of targeting *PCSK9*, underscoring the validity of our method for identifying novel drugs that could treat hyperlipidemia in the future. Other genes printed in black (e.g. *CELSR2*) in **Figure 4** are not targets of existing medications and therefore may be ideal for similar novel drug development.

### Replicating gene-statin interaction loci in independent samples of African-, Americas-, and European-like ancestry samples

After stratifying the AoU data^20^ into African-like (AFR), Americas-like (AMR), and European-like (EUR) ancestry groups, we conducted two replication analyses in the AoU data^20^: replication of the specific SNPs from the UKB analyses and replication within the LD blocks correcting for a multiple testing (p = 0.05/5 = 0.01) (**Tables 4 & 5**).

For replication of the specific SNPs, in the AoU EUR subsample, one SNP-statin interactions, and three other SNP-statin interactions showed a consistent, yet non-significant, pattern of associations (**Table 4**). In the AoU AFR ancestry group, one interaction replicated, but other SNPs had an unanticipated pattern of association. In the AMR ancestry group, two SNPs showed non-significant association with the expected pattern, and one SNP had an unanticipated pattern of association, and two SNPs were screened out of the analysis based on minor allele frequencies less than our minimum analytical threshold (MAF<0.01).

**Table 4:**
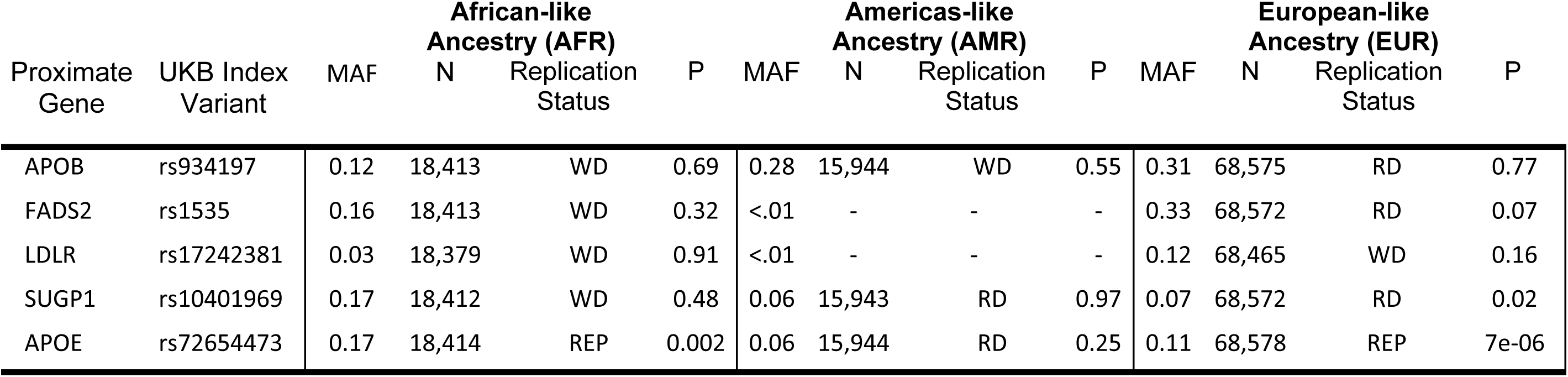
Comparison of the UK Biobank European gene-statin interaction lead variants with the lead variants in the African-like (AFR), Americas-like (AMR), and European-like (EUR) gene-statin GWAS analysis of LDL Cholesterol. MAF indicates the minor allele frequency in the AoU sample. N represents the sample size for the analysis of the specific SNP. For replication status, REP indicates that the pattern of marginal effects was in the right direction and statistically significant (p < 0.01); RD indicates that the pattern of marginal effects was in the right direction, but not significant (p > 0.01); and WD indicates that the pattern of marginal effects was in the wrong direction, regardless of statistical significance.

Second, we conducted a LD block replication analysis (**Table 5**). The most significant variant in the discovery sample may not be the lead variant in the replication sample due to differences in LD structure and allele frequencies across ancestry groups. For each of the five SNP-statin interaction loci, we analyzed variants within the associated LD blocks by selecting 1,000 proximate SNPs in each ancestry group (MAF > 0.01) centered around the most significant base pair position from the discovery analyses. Using a multiple testing significance threshold of p < 0.01, four of the five loci had at least one significant SNP in the EUR group, two of five loci in AMR group, and one of five in the AFR group. Importantly, the region that included *APOE* region showed significant replication across all three ancestry groups, highlighting the robustness of this gene-statin interaction across distinct populations.

**Table 5:**
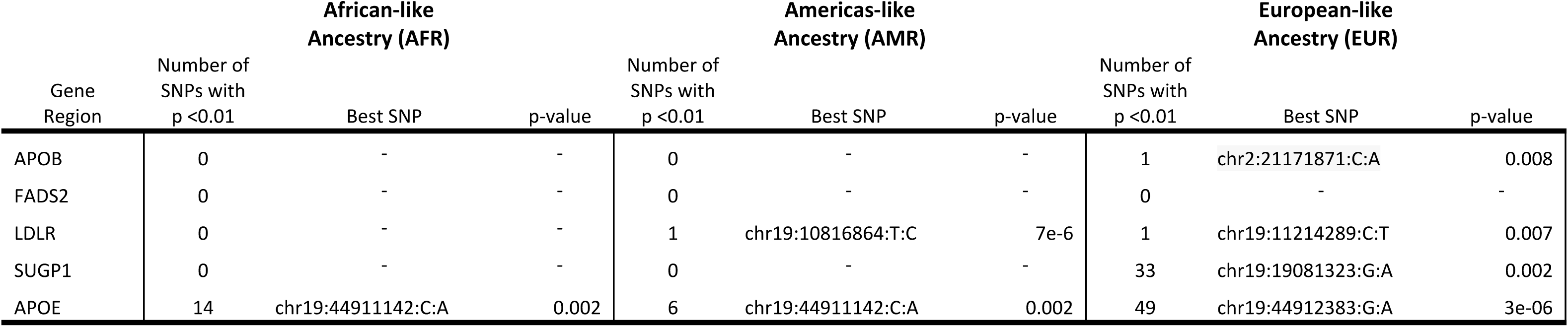
Regional Comparisons of 1,000 Variants centered around the lead variant from the discovery gene-statin interaction analysis in the African-like (AFR), Americas-like (AMR), and European-like (EUR) from the All of Us sample.

| Gene<br>Region | African-like<br>Ancestry (AFR) |  |  | Americas-like<br>Ancestry (AMR) |  |  | European-like<br>Ancestry (EUR) |  |  |
| --- | --- | --- | --- | --- | --- | --- | --- | --- | --- |
|  | Number of<br>SNPs with<br>p <0.01 | Best SNP | p-value | Number of<br>SNPs with<br>p <0.01 | Best SNP | p-value | Number of<br>SNPs with<br>p <0.01 | Best SNP | p-value |
| APOB | 0 | - | - | 0 | - | - | 1 | chr2:21171871:C:A | 0.008 |
| FADS2 | 0 | - | - | 0 | - | - | 0 | - | - |
| LDLR | 0 | - | - | 1 | chr19:10816864:T:C | 7e-6 | 1 | chr19:11214289:C:T | 0.007 |
| SUGP1 | 0 | - | - | 0 | - | - | 33 | chr19:19081323:G:A | 0.002 |
| APOE | 14 | chr19:44911142:C:A | 0.002 | 6 | chr19:44911142:C:A | 0.002 | 49 | chr19:44912383:G:A | 3e-06 |

## Discussion

Statins lower cholesterol for almost everyone, but statin responsiveness is a heterogeneous function of individual genetic differences. Despite phenotypic and genetic correlations among LDL-c, HDL-c, and TG, statins appear to specifically disrupt the genetic factors that contribute to the heritability of LDL-c. This finding is consistent with the clinical observation that statins lower the h^2^_SNP_ of LDL-c but leaves both HDL-c and TG relatively unchanged. In the discovery analyses, we identified five genome-wide significant gene-statin interactions that highlight genomic pathways that allow statins to lower LDL-c. Specifically, *SUGP1*/*MAU2* stimulates alternative splicing of *HMGCR,* decreasing its transcript stability and subsequently reducing cholesterol synthesis. The *SUGP1*/*MAU2*-statin interaction amplifies individual genetic differences in cholesterol synthesis reduction. Further, both *APOB* and *APOE* play pivotal roles in regulating lipid metabolism and homeostasis, notably in liver uptake and clearance via the LDL receptor, as coded for by *LDLR*^33,34^. Genetic variation in the *LDLR* gene is associated with elevated LDL-c, and statins lower LDL-c by upregulating *LDLR* expression thereby increasing the efficiency of the LDL receptor in liver cells ^35,36^. Accordingly, individual differences in *LDLR* may amplify LDL-c uptake in the liver^37,38^. Finally, *FADS2* plays a role in the regulation of cholesterol accumulation related to polyunsaturated fatty acids as the liver preferentially converts polyunsaturated fatty acids into ketone bodies rather than into LDL-c. The gene-statin interactions at these loci imply that statins will be more effective for individuals who have protective alleles in these regions. Taken together, the results reveal how statins dampen the genetic associations between these loci and LDL-c by interfering with multiple biological and genomic pathways that regulate cholesterol uptake, metabolism, and homeostasis.

In the AoU replication analyses, four of the five SNP-statin interactions from the UKB analysis had at least one significant interaction in the same LD block in the EUR sample. In the AFR and AMR replication samples, the results were a little mixed. Differences in allele frequencies across non-European ancestry groups reduce the power to detect both associations and interaction effects^39,40^. This obscures the distinction between ancestry-specific statin sensitivity and the lack of statistical power. The cross-ancestry replication of SNP-statin interactions in the *APOE* region, however, highlights the importance of the statin sensitivity for multi-ancestry populations.

Unfortunately, we were unable to replicate several previously published gene-statin interactions for LDL-c: specifically, variants in *SORT1*/*CELSR2*/*PSRC1*, *ABCG2*, *KIF6*, *LPA*, *CYP2C9*, and *SLCO1B1*^16^. While gene-statin interactions for these loci did not reach genome-wide significance, statin use reduced the absolute magnitude of the genetic associations for each locus. The lack of replication likely reflects several factors. First, our use of binary statin exposure without dosage information is better at detecting pharmacodynamic interactions with lipid pathways rather than pharmacokinetic genes like *CYP2C9*, and *SLCO1B1.* Second, these loci may have smaller interaction effect sizes than those we detected, requiring even larger samples. Additionally, prior studies used longitudinal or pre-post assessments, which focus on within-person changes in lipid values. This may provide greater sensitivity to detect interactions than our cross-sectional approach, which captures between-person differences that include additional sources of variation. However, longitudinal data tend to have markedly smaller sample sizes because they are more expensive to collect. As interaction methods require larger sample sizes than standard GWAS, we emphasized statistical power in our analyses. Furthermore, while the *Lp(a)*-statin interaction was not genome-wide significant for LDL-c, it was for TG, allowing for the possibility that the reduction in cardiovascular risk due to statin use may not be exclusively through LDL-c, validating the need and potential for new drug development for *Lp(a)*^41–43^. In contrast with previously reported gene-statin interactions, none of the loci we identified consist of single, lonely SNPs. Instead, we identified robust patterns of moderation across the LD region.

Our analyses identified 15 loci that remained genome-wide significant despite statin treatment. Many of these associations are near genes with well-validated links to LDL-c emphasizing that statins reduce but do not eliminate the genetic risk for hyperlipidemia^18^. Both the gene-statin interactions and the residual genetic associations have important pharmacogenomic implications. These residual genetic associations highlight potential statin-relevant future drug targets. Some of these loci are current statin targets and increasing the effectiveness of statins may further interrupt these genetic pathways. Specifically, variants in *APOB*, *APOE*/*PVRL2* and *LDLR*/*SMARCA4* remain significant in statin users, and therefore increasing statin efficacy may more efficiently interrupt these genetic pathways. Finally, we identified several genes that are not currently targeted by existing drugs (e.g., *LPA*). Existing evidence suggests novel drugs based on standard GWAS analyses are more likely to pass Stage 3 clinical trials than drugs developed using other methods^44–46^. Accordingly, just as standard GWAS have been effective for identifying genetic pathways that can be leveraged when developing pharmacological interventions^47^, the genomic pathways that remain significant for individuals who are already taking statins may be even more effective for drug development as they highlight treatment refractory genes.

Gene-statin interaction analyses provide pharmacogenomic insights from both disease-and patient-centered perspectives. From a disease-centered perspective, knowing which genes interact with statin use informs the biology of hyperlipidemia by identifying specific genomic pathways that statins interrupt. Many pharmacogenomic results have focused on the pharmacokinetic aspects of medication use (e.g., absorption, bioavailability, distribution, metabolization, and excretion). The broad implication of these findings is that poor metabolizers will require lower doses while ultrarapid metabolizers require higher doses to achieve optimal effects. Such findings are predicated on the central role of cytochrome P450 genes (e.g. CYP2D6 or *CYP3A4*). Because we did not examine dosage, we did not identify any genes from this class. Instead, our results highlight pharmacodynamic aspects of statin use (e.g., mechanisms of action with respect to genomic pathways), emphasizing a distinct perspective for personalized pharmacogenomic research. By further investigating how statins affect these pharmacodynamic genetic pathways and then integrating them with the existing pharmacokinetic results, it will become possible to identify individuals who are likely to benefit more (or less) from statin use based on their unique genotypes.

From a patient-centered perspective, our results highlight variants that are responsive to statin use, suggesting the possibility of tailoring treatment to individual patient’s genotypes. While statins decrease LDL-c by 33 mg/dL on average, each protective allele from *APOB*, *FADS2*, *LDLR*/*SMARCA4*, *SUGP1*/*MAU2*, and *APOE*/*PVRL2* provides an additional reduction (Figure 4). Knowing who will benefit the most from prescription medication, based on their genotype, has profound implications for prescribing medications, titrating dosage, and reducing the probability of adverse medication-induced side effects.

Our analytical methods provide a novel approach for assessing the genomic mechanisms of medication use broadly. We anticipate these methods will become increasingly applicable as high-quality genomic data continue to grow. The pharmacogenomic hypothesis is that medication will disrupt targeted genomic pathways, lower the phenotype heritability, or reduce or eliminate the genetic associations with variants in the targeted genes. Currently, however, medications are typically developed to target specific genetic pathways in drug naïve populations and tested with clinical trials. Because of this strategy, most current research excludes individuals who take statins, or other cholesterol-lowering medications. Confirming the efficacy of statins in a large general population sample of both users and non-users provides a unique opportunity to examine the pharmacogenomic sensitivity of the observed and intended gene targets. Our analyses simultaneously highlight genomic pathways that are altered by statin use, as well as those that remain significant after use. However, we refrain from making causal inferences based on these results, as respondents in population-based samples are not randomly assigned to statin use. Those who take statins likely have a higher genetic predisposition for elevated LDL-c as well as other cardiovascular outcomes (**Table 1**). Reciprocally, people with relatively low LDL-c are less likely to have genotypes that result in elevated cholesterol and thus have no need for statins (which they would not have been prescribed). As non-random statin treatment could potentially result in collider bias^48^, we conducted a simulation study (**Supplementary Information 15**) that demonstrated the genetic marginal effects are unaffected by non-random statin use but the statin use coefficient (β̂_2_ in Eq2) was underestimated. As such, the genetic marginal effects in this study are robust to the non-random prescription of statins for elevated cholesterol.

Statin pharmacogenomic studies have larger sample sizes than other gene-drug analyses, with the largest gene-statin pharmacogenomic sample consists of approximately 40,000 statin users^16^. Nevertheless, the current sample is 10x larger than the biggest existing statin sensitivity GWAS, because GWAS effect sizes are often very small, genome-wide gene-drug interaction analyses will require even larger sample sizes to identify pharmacogenomic loci. Existing evidence supports the fact that our novel gene-drug interaction methods can further refine our understanding of the biological mechanisms that regulate lipid production and metabolism, thereby improving treatment efficacy by focusing attention on the genetic signatures that are not minimized by statins.

To maximize the statistical power to detect SNP-statin interactions, we collapsed statins into a single, binary moderator, which has several implications. Because statin use was assessed via self-report in an unselected population sample, we assume that respondents answered accurately and patients were following the treatment regimens established with their physician, including compliance with the prescribed dose. Because medication use was based on self-reported data, we were confident in the respondent’s accuracy regarding the type of medications they used, while acknowledging that specific dosages had low levels of reliability. Because dosage information is particularly relevant for identifying pharmacokinetic effects, our analyses were not well-positioned to detect cytochrome P450 genes or *SLCO1B1*^1^. Nevertheless, we detected relevant pharmacodynamic processes related to gene-statin interactions. Further, and related to statistical power, by collapsing the five statins into a single dichotomous variable, we obscured the inherent pharmacological differences between statins. For example, atorvastatin is associated with a larger reduction in LDL-c levels compared with pravastatin^49^. While there are several commonly prescribed statins, in the current sample the majority of statin users were taking simvastatin followed by atorvastatin (**Table 1**). As such, our analyses primarily reflect interactions with these two medications. Future research would benefit from further refining these expectations by conducting gene-statin interaction analyses for specific statins. Because specific statins target slightly different biological pathways, future research should explore potential differential effects of each individual statin.

In summary, we demonstrated that statin use interferes with lipid levels by selectively reducing the heritability of LDL-c. We identified several loci that have genome-wide significant gene-statin interactions that provide insights into the biological mechanisms of action and suggest a process of determining which patients will benefit most from statin use. Furthermore, our analyses identified 15 loci that remained genome-wide significant despite treatment. Many of these associations are in genes with well-validated links to LDL-c implying that statins do not entirely disrupt genetic risk for high cholesterol, or that some genomic pathways are resistant to pharmacological intervention. These results are an important step towards using personal genomic information to provide targeted treatments to patients with hyperlipidemia.

## Methods

### Sample

For the discovery analyses, data come from the UK Biobank (UKB Application 57923). The full UKB dataset has approximately 500K UK individuals^50–52^. This sample is substantially larger and more representative of the UK general population than any other publicly available, genetically-informed dataset. For the discovery analyses, we restricted the sample to individuals of European ancestry for two reasons. First, as allele frequencies vary across ancestral populations, and if samples from multiple ancestral populations are analyzed jointly, population stratification could undermine the validity of the genetic associations^39,40^. Including ancestry PCs reduces, but does not entirely remove, the impact of population stratification^53^. Second, existing non-European sample sizes in the UKB do not provide sufficient power for genome-wide analyses^54,55^. This resulted in 387,420 European-like ancestry individuals.

Data for the replication analyses come from the AoU sample (November 17, 2025)^20^. The AoU data combine EHR data with genetic data for over 870,000 individuals. However, the heterogeneous nature of the data acquisition results in substantially lower sample sizes for most variables. We included individuals if they had valid measures of the key variables, including genotypes, LDL-c, and statins. This resulted in the inclusion of 69,123 European-like (EUR) ancestry individuals, 18,718 African-like (AFR) ancestry individuals, and 16,074 Americas-like (AMR) ancestry individuals.

### Outcome Variables

The UKB analyses use LDL-c, HDL-c, and TG measures from the initial data collection protocol. All lipid assays were conducted on blood samples, consistent with internationally recognized testing and calibration procedures. Lipids were initially measured in mmol/L and converted to mg/dL to simplify interpretation for American audiences. Specifically, we converted the values from mmol/l to mg/dl by multiplying the observed values for LDL and HDL by 38.67 and the observed values for TG by 88.57. Participants were not asked to fast prior to collecting the blood samples. Outcomes were treated as quantitative traits in the moderated GWAS analyses.

The AoU replication analyses focus on LDL-c. LDL-c measures were included if they could be converted into mg/dL without being extreme outliers (e.g. 1 mg/dL). As data come from available EHR entries, individuals can have multiple measures spanning years including LDL-c measures when they took statins and others where they did not. To take advantage of as much data as possible, for statin non-users we computed the individual’s mean LDL-c value across all valid measures. For statin users, we computed the mean of the LDL-c values within the window that the reported taking their statin prescription and excluding observations where the person had stopped taking statins.

### Moderator

There are numerous statins that are commonly prescribed for hyperlipidemia. In the UKB analyses, we focus on atorvastatin, fluvastatin, pravastatin, rosuvastatin, or simvastatin (or the brand-name equivalents). In the AoU analysis, we included the five statins used in the UKB analysis, as well as pitavastatin, lovastatin and cerivastatin. Patients were coded as statin users if they took any statin regardless of whether they took any other cholesterol-lowering medication (e.g., PCSK9 inhibitors) or used vitamin/dietary supplements to reduce cholesterol (e.g., niacin). Patients were coded as non-statin users if they only used vitamin/dietary supplements. Patients were excluded from all analyses if they ONLY used non-statin prescription medications to reduce their cholesterol.

### Genotypes

In the UKB data, DNA was extracted from blood samples taken from all willing participants. The DNA preparation and extraction was conducted by Affymetrix, and the initial quality control of the genomic data was conducted by the Wellcome Trust Centre for Human Genetics. Genotype imputation was conducted in IMPUTE2 using the UK10K as a reference panel. Imputed genotype data were used in all moderated GWAS analyses. Primary analyses were restricted to individuals of European ancestry to reduce the impact of population stratification on the results. To simplify the interpretation of the findings, we did not analyze insertion and deletion (indel) variants. For some respondents, individual SNPs did not pass quality control thresholds and were excluded from the analyses for those SNPs. This resulted in a sample size of 387,420. In the AoU data, short read whole genome sequencing data were generated from whole blood using the standardized pipeline across all Genome Centers to ensure standard QC methodologies and metrics, testing them with a series of validation experiments using previously characterized samples and commercially available reference standards^20,56^. This resulted in exceptionally high sensitivity and precision for common genomic variants.

### Covariates

To minimize the impact of known confounds of lipid variability, all the moderated GWAS analyses controlled for biological sex, age, the first 10 ancestry principal components (PCs), as well as interactions between the PCs and statins. The statin-PC interactions further account for the possibility of population stratification specific to the SNP-statin interaction coefficient that is not shared with the genetic main effects.

### Genome-wide gene-statin interaction analysis

All the moderated GWAS analyses of UKB data were conducted in R (v4.0)^57^ using the moderated GWAS functions in GW-SEM (v2.0) ^58,59^. The moderated GWAS analyses for the targeted replication in the AoU data were conducted in Python using equivalent techniques. Moderated GWAS analyses are an extension of standard GWAS where users conduct a series of regression analyses where a phenotype is regressed on each SNP in a genomic assay, adjusting for a set of predefined covariates such as age, sex, and the first 10 ancestry principal components to account for unobserved population stratification^60^. As such, the standard GWAS model for each variant is:

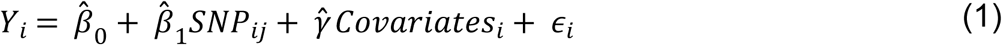

Where, in the current case, *Y*_*i*_ is the level of LDL-c, HDL-c, or TG for the i^th^ person, *SNP_ij_* is the j^th^ genetic variant for the i^th^ person, and *Covariates_i_* are the covariates for the i^th^ person (e.g., sex, age, and PCs). The estimate of the genetic association is β̂_1_, which is the most interesting parameter for most GWAS. The other estimates, β̂_0_ and γ̂, denote the intercept and regression coefficients for the covariates, respectively.

Genome-wide moderation analyses expand the standard GWAS model by regressing a phenotype on three independent variables: a) each SNP in the genomic assay, b) each individual’s statin use (the moderator), and the interaction between each SNP and the moderator, in addition to the covariates. To further rule out the possibility that population stratification may bias the interaction coefficients for the individual variants, interactions between the moderator and each of the first 10 ancestry principal components are also included as covariates^61^. Accordingly, the model for the genome-wide moderation analyses for each variant is:

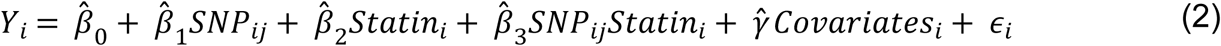

In the moderated GWAS model, both main effects, β̂_1_ and β̂_2_, depend on the interaction effect, β̂_3_. Importantly, β̂_3_ provides a test of whether the effect of a SNP on the phenotype changes at different levels of the environment. This parameter is difficult to interpret directly. Therefore, it is advantageous to calculate genetic marginal effects to examine how genetic association vary across levels of an environment.^60^

Finally, the interaction coefficient provides a formal test of the difference in the marginal genetic effects (for each locus) between the statin-takers and the non-statin-takers. Specifically, the interaction coefficient tells us whether the genetic associations differ in the statin vs non-statin groups. This coefficient must be interpreted with caution as interactions may amplify or diminish the main effect of the SNP on the phenotype (lipid) potentially leading to non-sensical conclusions. For example, if the interaction effect (β̂_3_) is in the opposite direction of the main effect (β̂_1_), it is possible to have a significant interaction coefficient despite not having significant genetic associations in either the statin or non-statin groups. Accordingly, we restrict our discussion of the significant interaction coefficients to those with meaningful substantive interpretations where the marginal effect is significant for at least the statin users or non-users.

### Marginal Genetic Effects

Summary statistics from the moderated GWAS of each of the lipids was used to calculate the marginal genetic effects and standard errors for the statin users and non-users. All calculations were done automatically in GW-SEM^59^. Genetic marginal effects are the association between a SNP and a phenotype at a specific level of an environment. In a standard GWAS model, the genetic marginal effect of the SNP is the regression coefficient (β̂_1_). Effectively, the standard GWAS regression coefficient is the effect of the SNP on the phenotype at the mean level of the moderating environment. In a moderated GWAS, the genetic marginal effect is a function of both genetic and moderating factors^60^. To calculate genetic marginal effects (β̂_*ME*_), we take the first derivative of the GxE GWAS model with respect to the SNP, leaving:

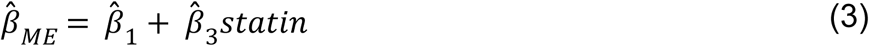

As respondents either took a statin (1) or did not (0), the marginal effect for the no statin group is simply β̂_1_ and the marginal effect for the statin group is β̂_1_ + β̂_3_. We can then calculate the standard errors of the genetic marginal effects (SE_ME_) using parameters from the variance-covariance (vcov) matrix of the moderated GWAS model for each SNP by:

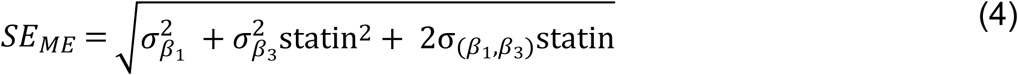

inserting the corresponding value of the statin group (0 or 1) that was used to calculate the genetic marginal effects. Note that for the no statin group, the SE reduces to 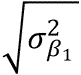 which is just the standard error of β̂_1_. The additional terms used to calculate the SE in the statin group incorporate any collinearity between the main effect and the interaction effect. After calculating the genetic marginal effect and the standard error, the z-statistic and p-value are easily calculated for use in subsequent analyses. This process is then repeated for each SNP that is analyzed. The process is automated in GW-SEM^59^, the only software platform that currently stores the σ_(β_1_,β_3_)_ statistic necessary to calculate the standard error of the marginal effects. To estimate group-specific SNP-heritability and genetic correlations, we applied LD score regression (LDSC) to the marginal effect summary statistics for statin users and non-users separately.

### Pharmacogenomic Analyses

The aim of the pharmacogenomic analyses was to collate our moderated GWAS results with pharmaceutical information regarding genetic targets for existing drugs using the pharmGKB.org database (the most extensive pharmacogenomic database available)^31^. We first extracted known gene targets for the five separate statins used as moderators in our analyses: atorvastatin, fluvastatin, pravastatin, rosuvastatin and simvastatin. The bottom edge of the triad of Figure 4 presents gene-targets. A full list of gene-targets is available in **Supplementary Table 6**. Second, we queried the pharmGKB.org database for drugs that are known to target the five genes with significant interaction effects: *APOB*, *FADS2*, *LDLR*/*SMARCA4, SUGP1*/*MAU2*, and *APOE*/*PVRL2*. The corresponding gene-drug combinations for the genes that interacted with statin use are presented in the left edge of the triad of Figure 4. Finally, we queried the pharmGKB.org database for drugs that target the 15 genes that remained significant for people taking statins. The corresponding gene-drug combinations for the residual genetic associations in the statin group are presented in the right edge of the triad in **Figure 4**.

## Declaration Statements

### Data Availability

Data were accessed through the UK Biobank Account Management System (application number 57923). In accordance with UK Biobank data access policies, raw can be accessed through the UK Biobank AMS after completing a data access agreement, but the sharing of raw phenotypic or genetic data is strictly prohibited. We gratefully acknowledge *All of Us* participants for their contributions, without whom this research would not have been possible. We also thank the National Institutes of Health’s *All of Us* Research Program for making available the participant data and samples examined in this study. The detailed summary statistics for the genome-wide significant loci are listing in supplementary tables. Portions of this research were conducted with the advanced computing resources provided by Texas A&M High Performance Research Computing. A complete set of summary statistics will be made available following the publication of the manuscript.

### Software Resources

**LDSC** (v1.0.1) https://github.com/bulik/ldsc**, GW-SEM** (v2.0) https://github.com/jpritikin/gwsem**, FUMA/MAGMA** (v 1.5.2) https://fuma.ctglab.nl, pharmGKB.org https://www.pharmgkb.org, **PLINK (v2),** https://www.cog-genomics.org/plink/2.0/.

## Acknowledgements

This work was supported by a Brain and Behavior Research Foundation BBRF 31397.

## Author Contributions

B.V. conceptualized the project, developed the methods, oversaw the analyses, and wrote the manuscript. J.H., A.M.A., and S.E.B. conducted the analyses. C.W.T, A.J.C, and J.M.H. edited the manuscript and provided scientific guidance.

## Competing Interests

The authors have no competing interests to declare.

## Notes

### Competing Interest Statement

The authors have declared no competing interest.

